# Mutation profiling of *KRAS* and *BRAF* in primary tumors and circulating tumor cells of colorectal cancer patients using PNA-LNA molecular switch

**DOI:** 10.1101/2025.06.01.657284

**Authors:** Md Sajedul Islam, Eliza Ranjit, Sharmin Aktar, Neda Moetamedirad, Cu Tai Lu, Muhammad J. A. Shiddiky, Vinod Gopalan, Alfred K. Lam

**Author notes:** Correspondence (M.J.A.S.), (V.G.), (A.K.L.).

## Abstract

Identifying *KRAS* and *BRAF* mutation status is essential for guiding targeted therapies and enhancing treatment outcomes in colorectal cancer (CRC). This study employs the “PNA-LNA molecular switch” to detect mutations in *KRAS* codon 12 (c.35G>T/G12V) and *BRAF* codon 600 (c.1799T>A/V600E) from primary tumours and circulating tumour cells (CTCs) in CRC patients, correlating mutation status with clinicopathological parameters. DNA was isolated from 71 primary tumours and 37 CTC samples. Mutation profiles were generated using the PNA-LNA molecular switch. *KRAS* mutations were detected in 26 primary tumours (36.6%) and 13 CTCs (26.8%), while *BRAF* mutations were observed in 19 primary tumours (26.8%) and 7 CTCs (19%). No significant correlation was observed between mutation status and clinicopathological parameters in primary tumours. However, *KRAS* G12V mutations in CTCs significantly correlated with lymph node metastasis (*p*=0.002), overall pathological stage (*p*=0.005), and lymphovascular invasion (*p*=0.034). *BRAF* V600E mutation status showed no significant clinicopathological associations. Validation of the PNA-LNA molecular switch against Next-Generation Sequencing (NGS) showed 89% concordance with *p*-values < 0.001 for both genes. This method is highly comparable to NGS for detecting *KRAS* and *BRAF* mutations and shows promise as a point-of-care diagnostic tool. Larger patient cohorts are required to confirm its clinical utility.

## 1. Introduction

According to the World Health Organization (WHO), colorectal cancer (CRC) ranks third in incidence and second in cancer-related mortality. The survival rate of CRC patients is strongly influenced by the stage at which the cancer is diagnosed[1]. Data from the American Cancer Society show that localized CRC has a 5-year survival rate of 90.5%, which drops sharply to 15% once distant metastasis occurs, emphasizing the critical importance of early detection[2].

Mutations in *KRAS*, *BRAF*, *PIK3CA*, and *P53* play a significant role in the development of CRC[3, 4]. Among these, *KRAS* and *BRAF*—two key oncogenes in the MAP-kinase signalling pathway—regulate essential cellular processes such as proliferation, differentiation, motility, and survival[5]. Somatic mutations in these genes disrupt normal cell cycle control, driving tumor progression[6]. *KRAS* mutations are more common than *BRAF* mutations in CRC, with prevalence rates of 53% and 10%, respectively[7, 8]. The most frequent *KRAS* mutations occur at codon 12 of exon 2 (e.g., c.35G>T/G12V and c.35G>A/G12D), while *BRAF* mutations typically occur at codon 600 of exon 15 (c.1799T>A/V600E)[9, 10]. These mutations are known to cause resistance to anti-EGFR therapies by persistently activating the MAP-kinase pathway, contributing to poor prognosis in patients with metastatic CRC[10-12]. As a result, routine testing for *KRAS* and *BRAF* mutations has become a standard component of CRC treatment planning.

In recent years, circulating tumor cells (CTCs) have emerged as promising biomarkers, offering a non-invasive way to explore tumor heterogeneity through genetic profiling[13, 14]. A study by Kalikaki et al. demonstrated that *KRAS* mutations found in CTC-enriched samples often reflect those in matched primary tumors[15]. However, the *KRAS* mutation status in CTCs can change throughout treatment, raising questions about their ability to consistently represent the tumour’s genetic landscape due to intratumor heterogeneity[16-18]. Despite this, profiling CTCs remains a valuable approach for monitoring tumor dynamics, although its clinical application for identifying *KRAS*/*BRAF* mutations and guiding immunotherapy still requires further validation.

Traditional mutation detection methods, such as polymerase chain reaction (PCR) and direct sequencing, are commonly used but come with drawbacks, including low sensitivity for detecting mutations at low allele frequencies, high costs, and lengthy processing times[19-23]. While next-generation sequencing (NGS) has become a clinical standard, it too remains costly and time-consuming. To overcome these challenges, we developed a novel technique that combines peptide nucleic acid (PNA) and locked nucleic acid (LNA) with loop-mediated isothermal amplification (LAMP)[20]. This method provides high sensitivity and specificity, enabling the detection of mutations in approximately 40 minutes at a lower cost compared to conventional approaches.

In this study, we employed the "PNA-LNA molecular switch" to detect *KRAS* and *BRAF* mutations in primary tumors and CTCs from CRC patients and examined their association with clinicopathological features. We also compared the results with NGS data to evaluate the clinical feasibility and sensitivity of this method.

## 2. Materials and Methods

### 2.1. Working principle

The PNA-LNA molecular switch identifies specific single-nucleotide polymorphisms (SNPs). PNAs are synthetic DNA analogs with repetitive units of N-(2-aminoethyl) glycine linked by amide bonds, replacing the phosphodiester backbone[24]. LNAs, by contrast, contain a methylene bridge between the 2’-oxygen and 4’-carbon atoms of the ribose, enhancing structural stability[25].

Two PNA probes targeting wild-type *KRAS* and *BRAF* genes and two LNA primers specific to *KRAS* (G12V) and *BRAF* (V600E) mutations were developed. PNAs bind to wild-type sequences during LAMP, inhibiting polymerase activity and blocking amplification. LNAs, on the contrary, bind to mutant sequences, enhancing amplification. Amplification is detected by measuring relative fluorescence units (RFU) above the threshold. Additionally, a colorimetric LAMP assay can identify mutations in the same experiment. Nucleotide incorporation lowers pH, turning phenol red from bright pink to yellow, while unamplified reactions remain pink. A yellow color change indicates target amplification, whereas pink indicates no amplification. **Figure 1** depicts the working principle of this study.

**Figure 1:**
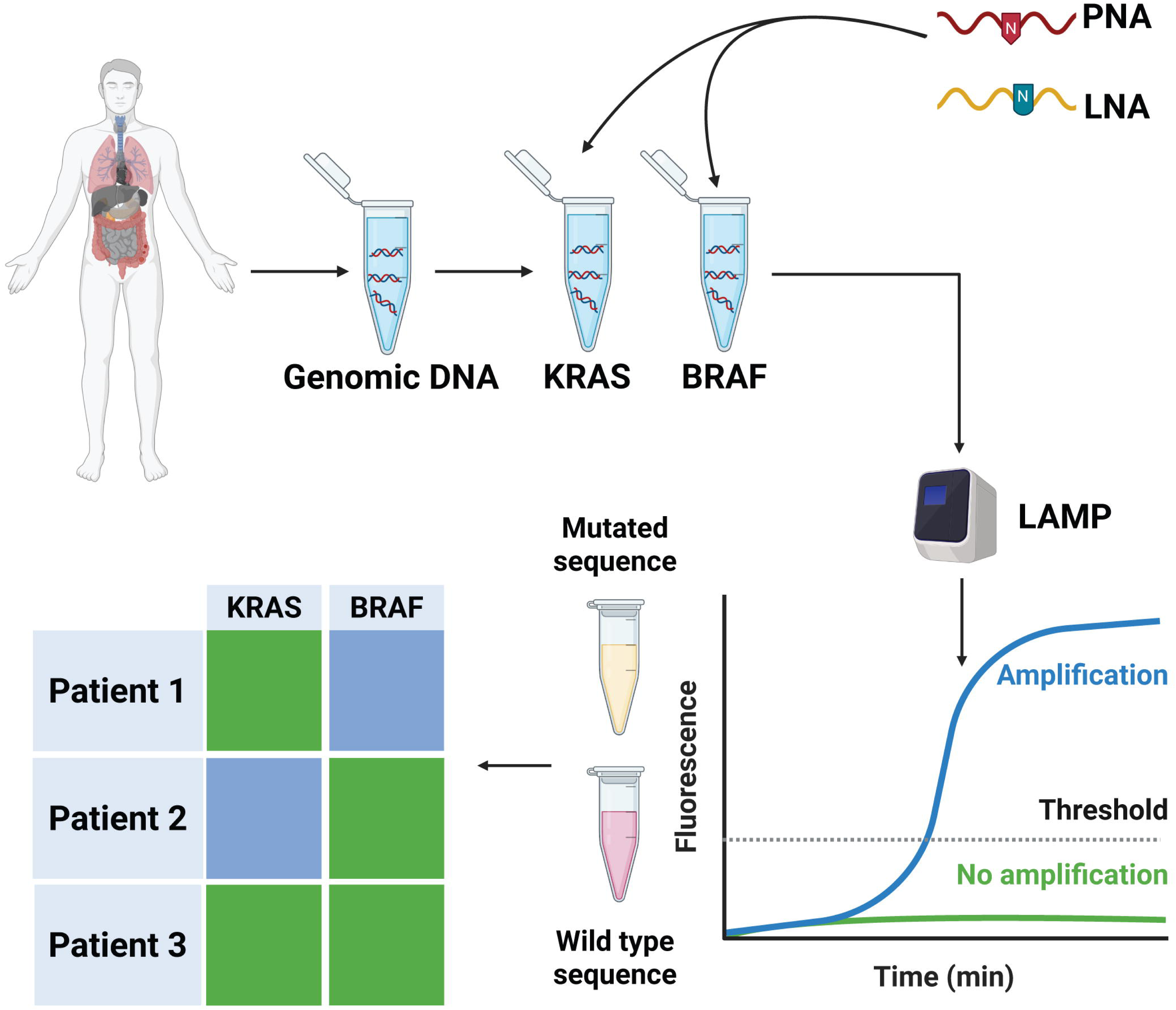
Working principle of PNA-LNA molecular switch. PNA probes specific for *KRAS* and *BRAF* genes bind to the wild-type sequence and block their amplification, while LNA primers enhance sequence amplification. Amplification can be visualized by fluorescence chromatogram and solution color change from pink to yellow. A mutation profile of the CRC is then created for *KRAS* and *BRAF* genes. This figure was generated in Biorender.

### 2.2. Patient recruitment

CRC tissue and peripheral blood samples were collected from patients undergoing colorectal carcinoma resections at Gold Coast University Hospital, Queensland, Australia. Tissue samples were snap-frozen in liquid nitrogen and stored at -80°C, while resected specimens were preserved in formalin for pathological analysis. Peripheral blood (10 ml) was collected in heparin tubes (Becton Dickinson) during surgery and processed within one hour to separate plasma, peripheral blood mononuclear cells (PBMCs), and enrich CTCs. Patient recruitment was unbiased, with ethical approval from the Griffith University Human Research Ethics Committee (GU Ref No: MSC/17/10/HREC). Written informed consent was obtained from each patient.

CRC tissues were analyzed for tumor size, site, histological subtype, microsatellite instability (MSI) status, lymphovascular invasion (LVI), and staging according to WHO criteria[26]. Cancer sites were classified as proximal (caecum, ascending, transverse colon) or distal (descending colon, sigmoid, rectum).

### 2.3. CTC enrichment and staining

CTCs were enriched using the EasySep™ Direct Human CTC Enrichment Kit (STEMCELL Technologies) following the manufacturer’s protocols. Detailed procedures were previously reported[27-29]. We then performed immunofluorescence staining to identify CTCs using four primary antibodies: mouse anti-EPCAM (Thermo Fisher Scientific), goat anti-SNAI1, mouse anti-E-cadherin, and goat anti-MMP9 (Santa Cruz Biotechnology). Secondary antibodies included rabbit anti-mouse IgG fluorescein isothiocyanate (FITC) and rabbit anti-goat IgG Texas Red (Sigma-Aldrich), along with Hoechst 33342 (Thermo Fisher Scientific) for nuclear staining. Stained cells were counted using a Nikon Ti-2 Widefield Microscope at 20× magnification. Cross-species binding of secondary antibodies was checked to prevent overlapping results, details of which were previously reported[27].

### 2.4. Extraction of DNA from tissues and CTCs

Following the manufacturer-provided protocols, we extracted the genomic DNA from the sectioned tissues and the CTCs using DNeasy Blood & Tissue kits (Qiagen) and AllPrep DNA/RNA mini kit, respectively. The concentration and purity of the extracted DNA were measured using a Nanodrop spectrophotometer (New England BioLabs). As the CTCs were scarce, whole-genome amplification was performed using the REPLI-g Advanced DNA Single Cell Kit (Qiagen) as previously described[29].

### 2.5. Primers and probe design

Two sets of four primary LAMP primers (FIP, BIP, F3, and B3) were designed for each of the *KRAS* and *BRAF* genes using PrimerExplorer v5 and NEB^®^ LAMP primer design tool. Additionally, two PNA probes were designed for each gene targeting the wild-type sequences, and two LNA primers were designed to target the mutated sequences (*KRAS* G12V and *BRAF* V600E) (**Table 1**). PNAs were purchased from PANAGENE, and the LNAs from IDT. The primers and probes were stored at -20L following resuspension in 1x Tris-EDTA buffer (IDT). The specificity of the primers and probes was verified using Primer Blast and UCSC in-silico PCR, ensuring no off-target binding to other oligonucleotide sequences.

**Table 1:**
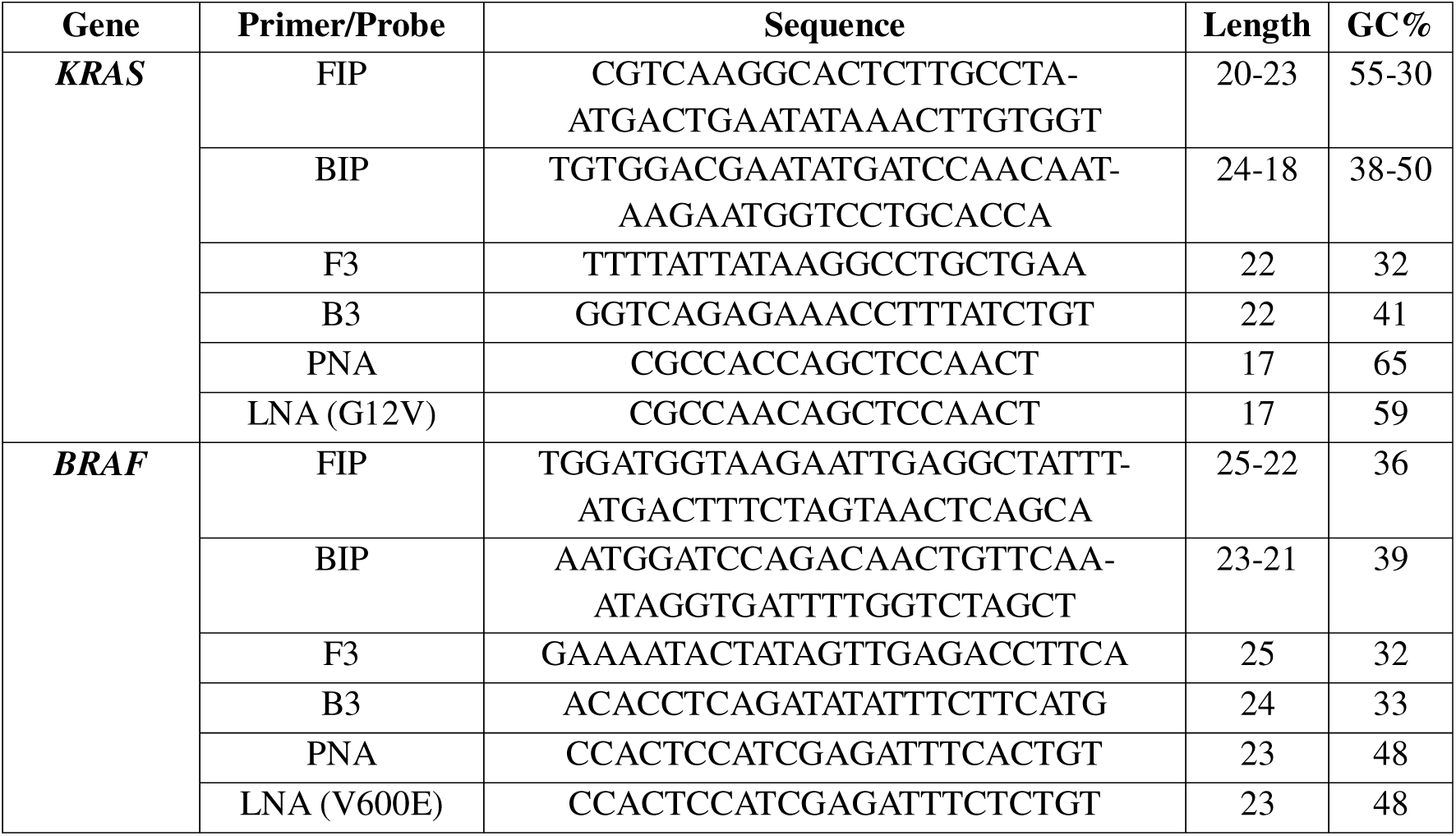
Primers and probe sequences.

### 2.6. LAMP reactions with PNA-LNA molecular switch

LAMP reactions were conducted using the WarmStart® Colorimetric and Fluorescent LAMP 2x Master Mix (NEB). The reaction conditions included primers and probes at the following concentrations: 1.6 μM FIP, 1.6 μM BIP, 0.2 μM F3, 0.2 μM B3, 1 μM LNA, and 0.5 μM PNA. The LAMP Fluorescent Dye (NEB) was used for each experiment alongside the colorimetric master mix in the same PCR tube. Each set of reactions included a positive control (excluding the PNA probe) and a negative control (excluding the DNA template). DNA amplification was performed using the QuantStudio™ 6 Flex Real-Time PCR system (Applied Biosystems). The reaction was initiated at 65°C for up to 90 minutes, followed by termination at 80°C for 2 minutes. Each reaction was performed in triplicate to ensure reproducibility. DNA amplification was measured by tracking changes in RFU over time, with the threshold time indicating when fluorescence intensity surpassed a set threshold. Positive samples that contained mutations or did not have any PNA changed color from pink to yellow, while negative samples remained pink. LAMP products were further analyzed using 1% agarose gel electrophoresis in 1x TAE buffer, stained with SYBR™ Safe DNA Gel Stain (Thermo Fisher Scientific), and visualized using a Gel documentation system (Bio-Rad).

### 2.7. Validation and statistical analysis

Results from the PNA-LNA molecular switch were compared with NGS variant analysis targeting *KRAS* (exons 2, 3, 4, and 5) and *BRAF* (exons 11 and 15) data from resected cancer tissues. The correlations between different parameters were studied by Fisher’s exact test or Pearson’s *chi-*square test using Statistical Package for Social Sciences (SPSS) version 29.0.2.0 (IBM). Additional statistical analyses and graphical representations were carried out using GraphPad Prism 10 and Microsoft Excel.

## 3. Results

### 3.1. Overall clinicopathological characteristics of the patients

In this study, we recruited 71 patients, consisting of 36 males and 35 females. Patients were selected without any sampling bias. The median age of the patients was 68, with an age range spanning from 28 to 92 years, of whom over 67% were above the age of 60. The tumor sizes of the large intestine adenocarcinomas ranged from 5 to 95 mm. Most of these tumors (63.4%) originated from the colon, and approximately 10% of these were mucinous adenocarcinomas. In terms of grading, 50% of the tumors were classified as grade 2 adenocarcinomas. Notably, almost 70% of the adenocarcinomas were either stage III or IV (advanced stages), and around half of the cases showed positive lymph node involvement, indicating carcinoma infiltration into the lymph nodes. Despite the high proportion of advanced-stage tumors, the overall pathological stage distribution between early-stage and advanced-stage adenocarcinomas was relatively balanced. Furthermore, around 40% of the patients had positive LVI status, whereas 22.5% had high MSI. Among the 71 patients, CTCs were detected in 37 patients, with counts ranging from 2 to 200. **Table 2** summarises the demographic and clinicopathological characteristics of the recruited patients.

**Table 2:** Demographic characteristics of the recruited patients with colorectal adenocarcinoma.

| Demographic characteristics | Tissue (n = 71) | CTC (n = 37) |
| --- | --- | --- |
| **Gender** |  |  |
| Male | 36 (50.7%) | 17 (45.9%) |
| Female | 35 (49.3%) | 20 (54.1%) |
| **Age** |  |  |
| ≤ 60 years | 23 (32.4%) | 13 (35.1%) |
| > 60 years | 48 (67.6%) | 24 (64.9%) |
| **Size** |  |  |
| ≤ 40 mm | 41 (57.7%) | 23 (62.2%) |
| > 40 mm | 30 (42.3%) | 14 (37.8%) |
| **Site** |  |  |
| Colon | 45 (63.4%) | 21 (56.8%) |
| Rectum | 26 (36.6%) | 16 (43.2%) |
| **Subtype of adenocarcinoma** |  |  |
| Classic | 64 (90.1%) | 35 (94.6%) |
| Mucinous | 7 (9.9%) | 2 (5.4%) |
| **Grade** |  |  |
| Well (1) | 13 (18.3%) | 6 (16.2%) |
| Moderate (2) | 50 (70.4%) | 27 (73%) |
| Poor (3) | 8 (11.3%) | 4 (10.8%) |
| **T Stage** |  |  |
| I or II | 22 (31%) | 13 (35.1%) |
| III or IV | 49 (69%) | 24 (64.9%) |
| **Lymph Node Status** |  |  |
| Negative | 36 (50.7%) | 19 (51.4%) |
| Positive | 35 (49.3%) | 18 (48.4%) |
| **Distant metastasis** |  |  |
| Negative | 60 (84.5%) | 32 (86.5%) |
| Positive | 11 (15.5%) | 5 (13.5%) |
| **Overall Pathological Stage** |  |  |
| I or II | 34 (47.9%) | 18 (48.6%) |
| III or IV | 37 (52.1%) | 19 (51.4%) |
| **Perineural invasion** |  |  |
| Negative | 61 (85.9%) | 32 (86.5%) |
| Positive | 10 (14.1%) | 5 (13.5%) |
| **Lymphovascular invasion** |  |  |
| Negative | 43 (60.6%) | 21 (56.8%) |
| Positive | 28 (39.4%) | 16 (43.2%) |
| **Microsatellite instability** |  |  |
| Stable | 55 (77.5%) | 29 (78.4%) |
| High | 16 (22.5%) | 8 (21.6%) |

### 3.2. Assay optimization and analytical performance

In our previous work, we detailed the optimization and analytical performance of this assay[20]. Briefly, we performed serial dilutions of synthetic DNA targeting the gene sequence, ranging from 10 pg/μl (10^7^ DNA copies/μl) to 1 ag/μl (1 DNA copy/μl). The optimal concentration for subsequent analysis was determined to be 1 fg/μl (10^3^ DNA copies/μl), as this yielded an amplification threshold time of under 40 minutes. We further optimized the PNA probe and LNA primer concentrations to 0.5 μM and 1 μM, respectively. For clinical samples, including tissue and CTCs, we used a DNA concentration of 1 ng/μl (10^3^ DNA copies/μl).

Specificity testing with synthetic and cell line DNA showed selective amplification of the mutated sequences, with no amplification of wild-type sequences or non-targets. The method also successfully distinguished between different mutant subtypes at the same locus (*KRAS* G12V vs G12D) within approximately 10 minutes. In terms of sensitivity, the assay demonstrated linearity, with amplification signals directly proportional to the DNA input. The assay maintained this relationship across the dynamic range for synthetic target DNA, yielding an excellent efficiency, as indicated by an R² value of 0.94. The limit of detection (LOD) was determined to be 0.00009, corresponding to a minimum detectable concentration of 1 ag/μl (1 DNA copy/μl).

### 3.3. *KRAS* mutation frequency

Among 71 adenocarcinomas, 26 (36.6%) harboured G12V mutation in the *KRAS* codon 12. Of the 37 patients who tested positive for CTCs, 13 cases (26.8%) exhibited the *KRAS* G12V mutation within their CTCs. After analyzing the mutations in both the tissue and CTC samples, 18 cases (49%) were wild-type, and 11 cases (30%) were mutant in both the tumors and corresponding CTC samples. Interestingly, two cases showed acquired mutations in codon 12 of the *KRAS* gene within the CTCs, despite being wild-type in the primary tumor tissue. Conversely, in six cases, the CTCs were wild-type, while the corresponding tissues carried the *KRAS* mutation (*p*<0.001) (**Table 3**).

**Table 3:**
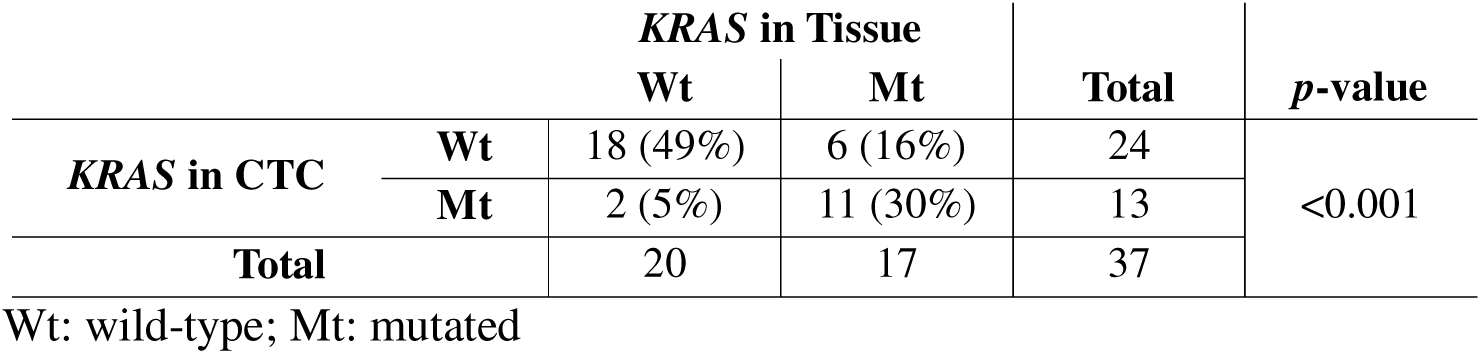
Mutation status of *KRAS* (G12V) from tissue and CTC.

Comparing the results of 27 adenocarcinoma cases obtained through the PNA-LNA molecular switch with those from NGS analysis, 17 cases were consistently identified as wild-type, and 7 were identified as mutant for *KRAS* G12V by both methods. There were two cases where our method detected mutations that NGS failed to identify. On the other hand, there was one case where NGS detected a mutation that the PNA-LNA molecular switch did not identify (*p*<0.001) (**Table 4**).

**Table 4:** Correlation of *KRAS* mutation status between PNA-LNA molecular switch and NGS.

|  |  | ***KRAS* in Tissue by Sequencing** |  | **Total** | ***p*-value** |
| --- | --- | --- | --- | --- | --- |
|  |  | **Wt** | **Mt** |  |  |
| ***KRAS* in Tissue by LAMP** | **Wt** | 17 (63%) | 2 (7%) | 19 | <0.001 |
|  | **Mt** | 1 (4%) | 7 (26%) | 8 |  |
| **Total** |  | 18 | 9 | 27 |  |

Wt: wild-type; Mt: mutated

### 3.4. *BRAF* mutation frequency

For the *BRAF* gene analysis, 19 cases (26.8%) were found to harbour the *BRAF* V600E mutation. Among the 37 CTC-positive patients, 27 (73%) were wild-type, and 7 cases (19%) carried the *BRAF* V600E mutation in both the tissues and CTCs. In alignment with previous observations, two cases exhibited an acquired *BRAF* mutation in the CTCs, even though the primary adenocarcinoma was wild-type. Conversely, one patient exhibited the *BRAF* V600E mutation in the adenocarcinoma but not in the corresponding CTC (*p*<0.001) (**Table 5**). Among the 27 cases analyzed with both the PNA-LNA molecular switch and NGS, 18 cases were consistently wild-type, while 6 cases were *BRAF* mutant in both methods. However, 2 cases showed *BRAF* mutations with the PNA-LNA molecular switch that were not detected by sequencing, and in one case, sequencing identified a *BRAF* mutation that was not detected by our method (*p*<0.001) (**Table 6**).

**Table 5:**
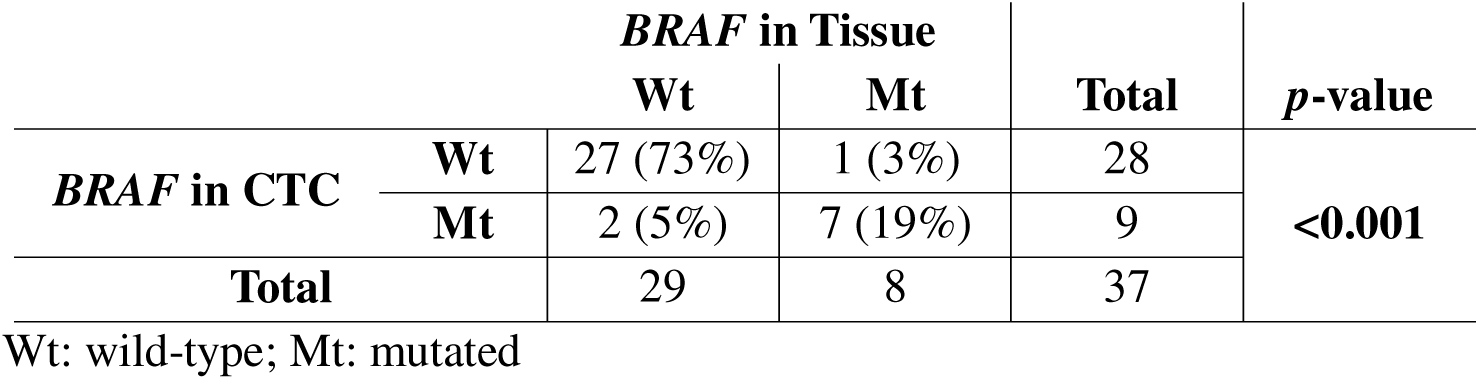
Mutation status of *BRAF* (V600E) from tissue and CTC.

|  |  | ***BRAF* in Tissue** |  | **Total** | ***p*-value** |
| --- | --- | --- | --- | --- | --- |
|  |  | **Wt** | **Mt** |  |  |
| ***BRAF* in CTC** | **Wt** | 27 (73%) | 1 (3%) | 28 | <0.001 |
|  | **Mt** | 2 (5%) | 7 (19%) | 9 |  |
| **Total** |  | 29 | 8 | 37 |  |

Wt: wild-type; Mt: mutated

**Table 6:** Correlation of *BRAF* mutation status between PNA-LNA molecular switch and NGS.

|  |  | ***BRAF* in Tissue by Sequencing** |  | **Total** | ***p*-value** |
| --- | --- | --- | --- | --- | --- |
|  |  | **Wt** | **Mt** |  |  |
| ***BRAF* in Tissue by LAMP** | **Wt** | 18 (67%) | 2 (7%) | 20 | <0.001 |
|  | **Mt** | 1 (4%) | 6 (22%) | 7 |  |
| **Total** |  | 19 | 8 | 27 |  |

Wt: wild-type; Mt: mutated

Figure 2 presents an amplification chromatogram with representative LAMP results obtained from the PNA-LNA molecular switch. **Supplementary Table S1** provides the mutation status of each patient for *KRAS* (G12V) and *BRAF* (V600E) from primary adenocarcinoma and matched CTCs, along with the NGS data.

**Figure 2:**
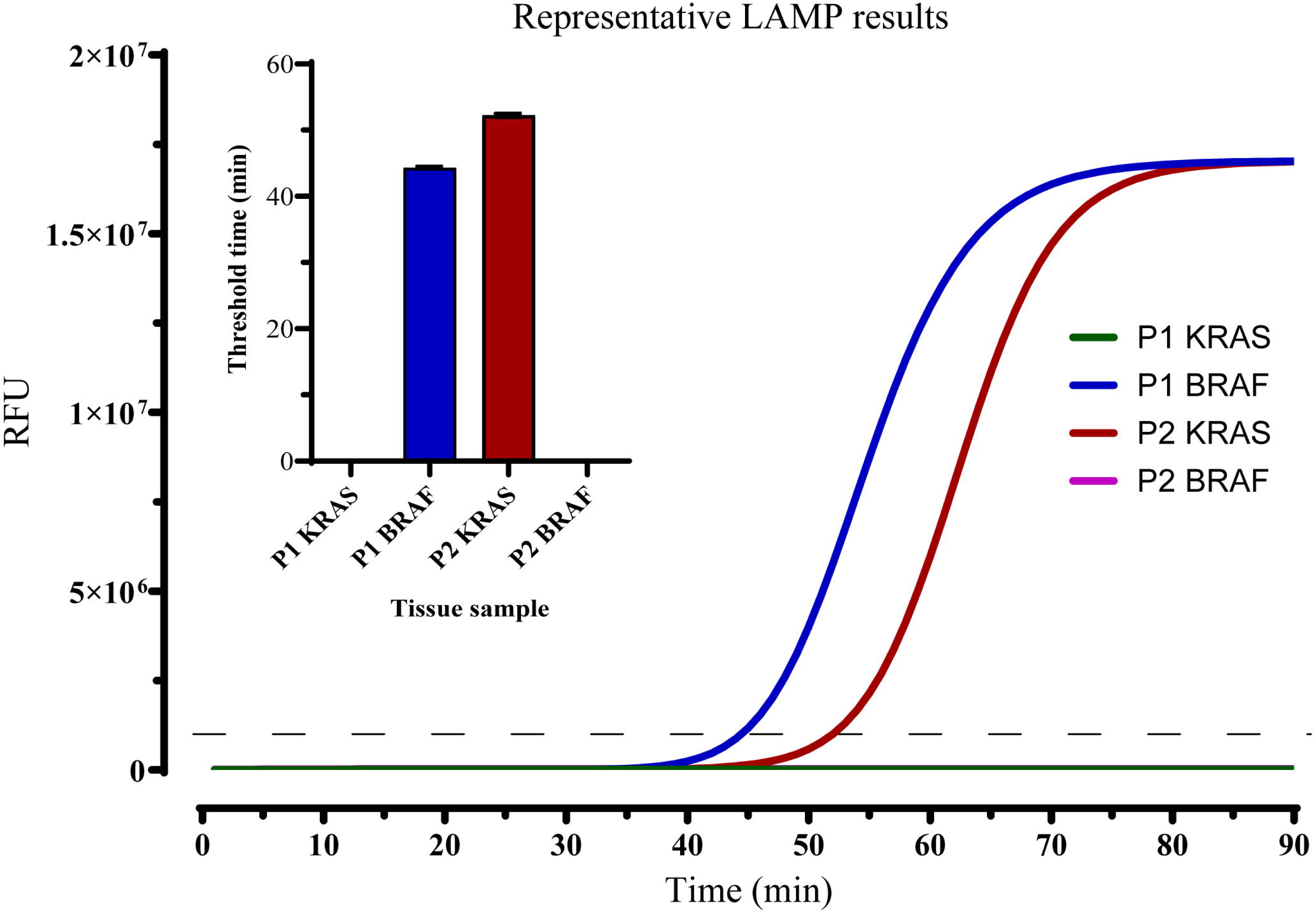
Representative chromatogram for LAMP results. Amplified chromatograms above the threshold level indicate the presence of mutant sequences, while chromatograms for wild-type sequences remain below the threshold. The inset shows the corresponding amplification threshold time bar diagram for each of the experiments, highlighting the differences in amplification times between wild-type and mutant sequences.

### 3.5. *KRAS* mutation and clinicopathological correlations

**Table 7** represents the correlation between *KRAS* (G12V) mutations found in primary tumors and corresponding CTCs with various clinicopathological characteristics in patients with CRC. The presence of *KRAS* mutation from the primary adenocarcinoma did not show significant associations with any clinicopathological factors. Nevertheless, a higher prevalence of *KRAS* mutations was observed in carcinomas with advanced local spread (T stage) (*p*=0.607) and higher overall pathological stage (*p*=0.138), although these trends were not statistically significant. Also, carcinomas exhibiting an MSI phenotype showed a lower frequency of *KRAS* mutations (*p*=0.380), and patients with perineural invasion were less likely to harbor *KRAS* mutations (*p*=0.155), though neither finding reached statistical significance.

**Table 7:** Correlations of *KRAS* mutation (G12V) with clinicopathological characteristics.

| Characteristics | Tissue |  |  | CTC |  |  |
| --- | --- | --- | --- | --- | --- | --- |
|  | Wt (%) | Mt (%) | *p*-value | Wt (%) | Mt (%) | *p*-value |
| **Gender** |  |  |  |  |  |  |
| Male | 24 (67%) | 12 (33%) | 0.627 | 10 (59%) | 7 (41%) | 0.512 |
| Female | 21 (67%) | 14 (33%) |  | 14 (70%) | 6 (30%) |  |
| **Age** |  |  |  |  |  |  |
| ≤ 60 years | 10 (43%) | 13 (57%) | 0.423 | 5 (38%) | 8 (62%) | 0.097 |
| > 60 years | 35 (73%) | 13 (27%) |  | 19 (79%) | 5 (21%) |  |
| **Size** |  |  |  |  |  |  |
| ≤ 40 mm | 23 (56%) | 18 (44%) | 0.325 | 15 (65%) | 8 (35%) | 0.391 |
| > 40 mm | 22 (73%) | 8 (27%) |  | 9 (64%) | 5 (36%) |  |
| **Site** |  |  |  |  |  |  |
| Colon | 30 (67%) | 15 (33%) | 0.458 | 15 (71%) | 6 (29%) | 0.489 |
| Rectum | 15 (58%) | 11 (42%) |  | 9 (56%) | 7 (44%) |  |
| **Subtype of adenocarcinoma** |  |  |  |  |  |  |
| Classic | 41 (64%) | 23 (36%) | 0.701 | 23 (66%) | 12 (34%) | 1.00 |
| Mucinous | 4 (57%) | 3 (43%) |  | 1 (50%) | 1 (50%) |  |
| **Grade** |  |  |  |  |  |  |
| Well (1) | 9 (69%) | 4 (31%) | 0.889 | 5 (83%) | 1 (17%) | 0.602 |
| Moderate (2) | 31 (62%) | 19 (38%) |  | 17 (63%) | 10 (37%) |  |
| Poor (3) | 5 (62%) | 3 (38%) |  | 2 (50%) | 2 (50%) |  |
| **T Stage** |  |  |  |  |  |  |
| I or II | 15 (68%) | 7 (32%) | 0.607 | 10 (77%) | 3 (23%) | 0.305 |
| III or IV | 30 (61%) | 19 (39%) |  | 14 (58%) | 10 (42%) |  |
| **Lymph Node Status** |  |  |  |  |  |  |
| Negative | 25 (69%) | 11 (31%) | 0.330 | 17 (89%) | 2 (11%) | **0.002** |
| Positive | 20 (57%) | 15 (43%) |  | 7 (38%) | 11 (62%) |  |
| **Distant Metastasis** |  |  |  |  |  |  |
| Negative | 38 (63%) | 22 (27%) | 1.00 | 21 (66%) | 11 (34%) | 1.00 |
| Positive | 7 (63%) | 4 (27%) |  | 3 (60%) | 2 (40%) |  |
| **Overall Pathological Stage** |  |  |  |  |  |  |
| I or II | 25 (73%) | 9 (27%) | 0.138 | 16 (89%) | 2 (11%) | **0.005** |
| III or IV | 20 (54%) | 17 (46%) |  | 8 (42%) | 11 (58%) |  |
| **Perineural invasion** |  |  |  |  |  |  |
| Negative | 41 (67%) | 20 (33%) | 0.155 | 22 (69%) | 10 (31%) | 0.321 |
| Positive | 4 (40%) | 6 (60%) |  | 2 (40%) | 3 (60%) |  |
| **Lymphovascular invasion** |  |  |  |  |  |  |
| Negative | 30 (70%) | 13 (30%) | 0.210 | 17 (81%) | 4 (19%) | **0.036** |
| Positive | 15 (54%) | 13 (46%) |  | 7 (44%) | 9 (56%) |  |
| **Microsatellite instability** |  |  |  |  |  |  |
| Stable | 33 (60%) | 22 (40%) | 0.380 | 17 (59%) | 12 (41%) | 0.216 |
| High | 12 (75%) | 4 (25%) |  | 7 (88%) | 1 (12%) |  |

On the other hand, *KRAS* (G12V) mutations in CTCs correlated significantly with clinicopathological characteristics like lymph node metastasis (*p*=0.002), overall pathological stage (*p*=0.005), and lymphovascular invasion (*p*=0.036). While not statistically significant, *KRAS* mutations in CTCs were also more common in carcinomas with advanced local spread (*p*=0.305). Similarly, high MSI in carcinoma was associated with fewer *KRAS* mutations in CTCs (*p*=0.216). These findings suggest that while *KRAS* mutations in primary tumors may not exhibit strong clinicopathological associations, mutations in CTCs may have potential clinical relevance, particularly with lymph node involvement, pathological staging, and LVI in CRC patients.

### 3.6. *BRAF* mutation and clinicopathological correlation in colorectal adenocarcinoma

No significant associations were found between *BRAF* mutations and the clinical pathological parameters, including tumour stage, MSI, or perineural invasion. Despite the lack of statistical significance, mutations in the *BRAF* gene were more prevalent in advanced local spread in primary tumors (*p*=0.387) but not in CTCs. Similarly, patients with high MSI were associated with fewer *BRAF* mutations in both primary adenocarcinomas (*p*=0.110) and CTCs (*p*=0.078), though these associations did not reach statistical significance. Moreover, carcinomas with perineural invasion also showed a tendency for fewer *BRAF* mutations in primary adenocarcinoma (*p*=0.719) and no mutations in CTCs (*p*=0.307). Overall, the distribution of *BRAF* mutation status was relatively even across clinical parameters, with no significant correlations observed. This suggests that *BRAF* mutations may not be closely linked with the clinicopathological features of CRC in this cohort.

**Table 8** summarises the correlation between *BRAF* (V600E) mutations in primary adenocarcinomas and corresponding CTCs with various clinicopathological characteristics in CRC patients.

**Table 8:** Correlation of *BRAF* mutation (V600E) with clinicopathological characteristics.

|  | Tissue |  |  | CTC |  |  |
| --- | --- | --- | --- | --- | --- | --- |
| Characteristics | Wt (%) | Mt (%) | *p*-value | Wt (%) | Mt (%) | *p*-value |
| **Gender** |  |  |  |  |  |  |
| Male | 25 (69%) | 10 (31%) | 0.793 | 13 (76%) | 4 (24%) | 1.00 |
| Female | 27 (77%) | 9 (23%) |  | 15 (75%) | 5 (25%) |  |
| **Age** |  |  |  |  |  |  |
| ≤ 60 years | 21 (91%) | 2 (9%) | 0.346 | 11 (85%) | 2 (15%) | 0.568 |
| > 60 years | 31 (65%) | 17 (35%) |  | 17 (71%) | 7 (29%) |  |
| **Size** |  |  |  |  |  |  |
| ≤ 40 mm | 33 (80%) | 8 (20%) | 0.291 | 20 (87%) | 3 (13%) | 0.572 |
| > 40 mm | 19 (63%) | 11 (37%) |  | 8 (57%) | 6 (43%) |  |
| **Site** |  |  |  |  |  |  |
| Colon | 32 (71%) | 13 (29%) | 0.782 | 15 (71%) | 6 (29%) | 0.702 |
| Rectum | 20 (77%) | 6 (23%) |  | 13 (81%) | 3 (19%) |  |
| **Subtype of adenocarcinoma** |  |  |  |  |  |  |
| Classic | 47 (73%) | 17 (27%) | 1.00 | 26 (74%) | 9 (26%) | 1.00 |
| Mucinous | 5 (71%) | 2 (29%) |  | 2 (100%) | 0 (0%) |  |
| **Grade** |  |  |  |  |  |  |
| Well (1) | 10 (77%) | 3 (23%) | 0.288 | 3 (50%) | 3 (50%) | 0.267 |
| Moderate (2) | 38 (76%) | 12 (24%) |  | 22 (81%) | 5 (19%) |  |
| Poor (3) | 4 (50%) | 4 (50%) |  | 3 (75%) | 1 (25%) |  |
| **T Stage** |  |  |  |  |  |  |
| I or II | 18 (81%) | 4 (19%) | 0.387 | 9 (69%) | 4 (31%) | 0.691 |
| III or IV | 34 (69%) | 15 (31%) |  | 19 (79%) | 5 (21%) |  |
| **Lymph Node Status** |  |  |  |  |  |  |
| Negative | 25 (69%) | 11 (31%) | 0.594 | 13 (68%) | 6 (32%) | 0.447 |
| Positive | 27 (77%) | 8 (23%) |  | 15 (83%) | 3 (17%) |  |
| **Distant Metastasis** |  |  |  |  |  |  |
| Negative | 42 (70%) | 18 (30%) | 0.267 | 23 (72%) | 9 (28%) | 0.307 |
| Positive | 10 (90%) | 1 (10%) |  | 5 (100%) | 0 (0%) |  |
| **Overall Pathological Stage** |  |  |  |  |  |  |
| I or II | 23 (68%) | 11 (32%) | 0.422 | 12 (67%) | 6 (33%) | 0.269 |
| III or IV | 29 (78%) | 8 (22%) |  | 16 (84%) | 3 (16%) |  |
| **Perineural invasion** |  |  |  |  |  |  |
| Negative | 44 (72%) | 17 (28%) | 0.719 | 23 (72%) | 9 (28%) | 0.307 |
| Positive | 8 (80%) | 2 (20%) |  | 5 (100%) | 0 (0%) |  |
| **Lymphovascular invasion** |  |  |  |  |  |  |
| Negative | 30 (70%) | 13 (30%) | 0.584 | 14 (67%) | 7 (33%) | 0.248 |
| Positive | 22 (79%) | 6 (21%) |  | 14 (88%) | 2 (12%) |  |
| **Microsatellite instability** |  |  |  |  |  |  |
| Stable | 43 (78%) | 12 (22%) | 0.110 | 24 (83%) | 5 (17%) | 0.078 |
| High | 9 (56%) | 7 (44%) |  | 4 (50%) | 4 (50%) |  |

## Discussion

The PNA-LNA molecular switch integrates synthetic DNA analogues, such as PNA and LNA, with LAMP to enable rapid and cost-effective detection of specific point mutations. The method demonstrates high sensitivity (detecting 1 DNA copy/µL), specificity, and reproducibility, and effectively differentiates between mutant and wild-type sequences, as well as identifying low allelic frequency mutations. Compared to traditional methods like Sanger sequencing, this assay is faster (approximately 40 minutes for clinical samples), more affordable (approximately A$7.50 per experiment), and simpler, requiring minimal equipment. In this study, we utilized the PNA-LNA molecular switch to assess the mutation status of *KRAS* G12V and *BRAF* V600E in primary tumor tissues and CTCs from colorectal cancer patients. The results obtained from this technique were compared with the corresponding NGS data. Our findings highlight the potential of the PNA-LNA molecular switch for efficient, cost-effective mutation analysis, offering valuable utility in the clinical management of CRC. A brief comparison of the analytical performance of this assay with contemporary methods is provided in **Table 9**.

**Table 9:**
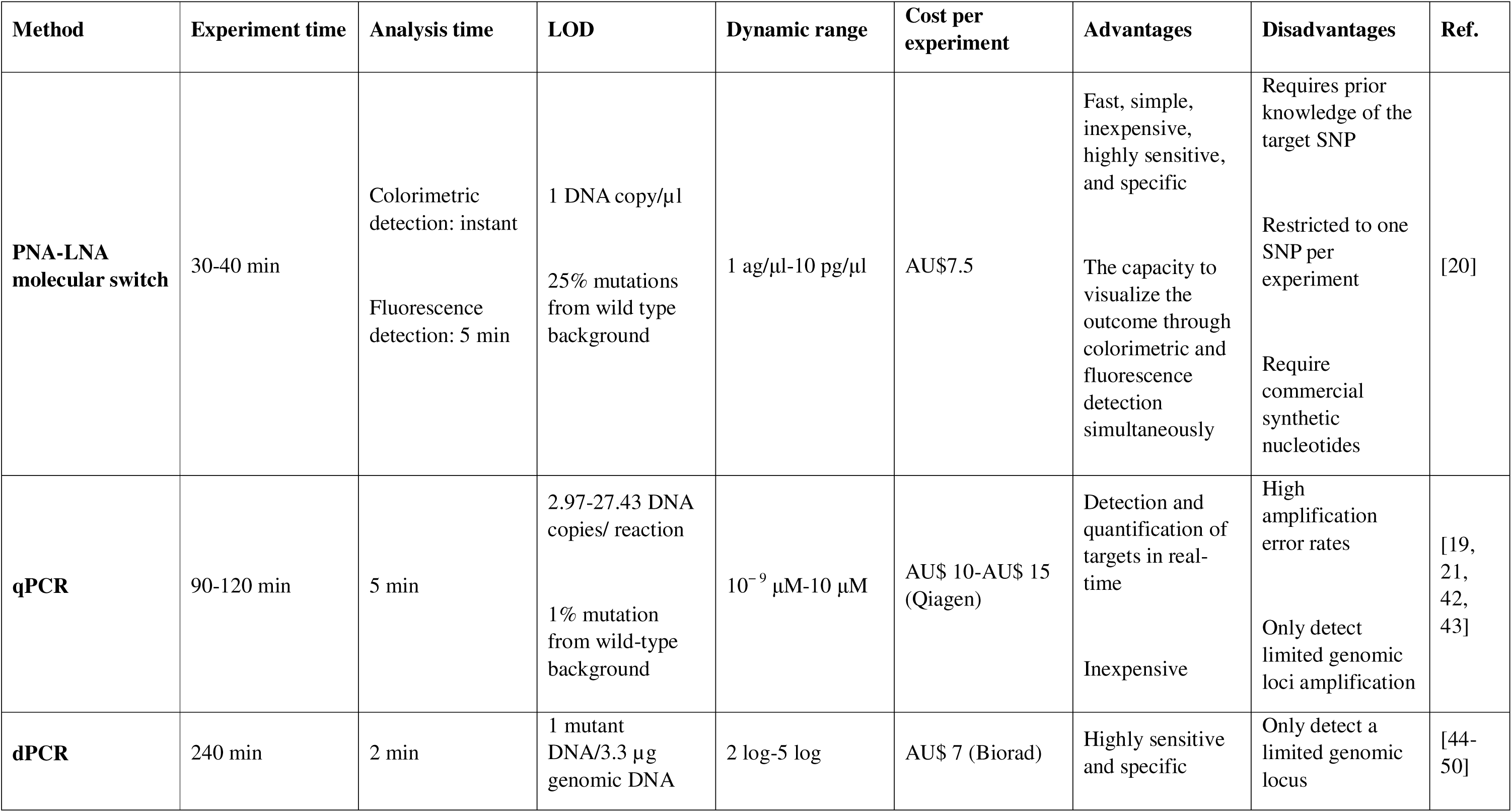
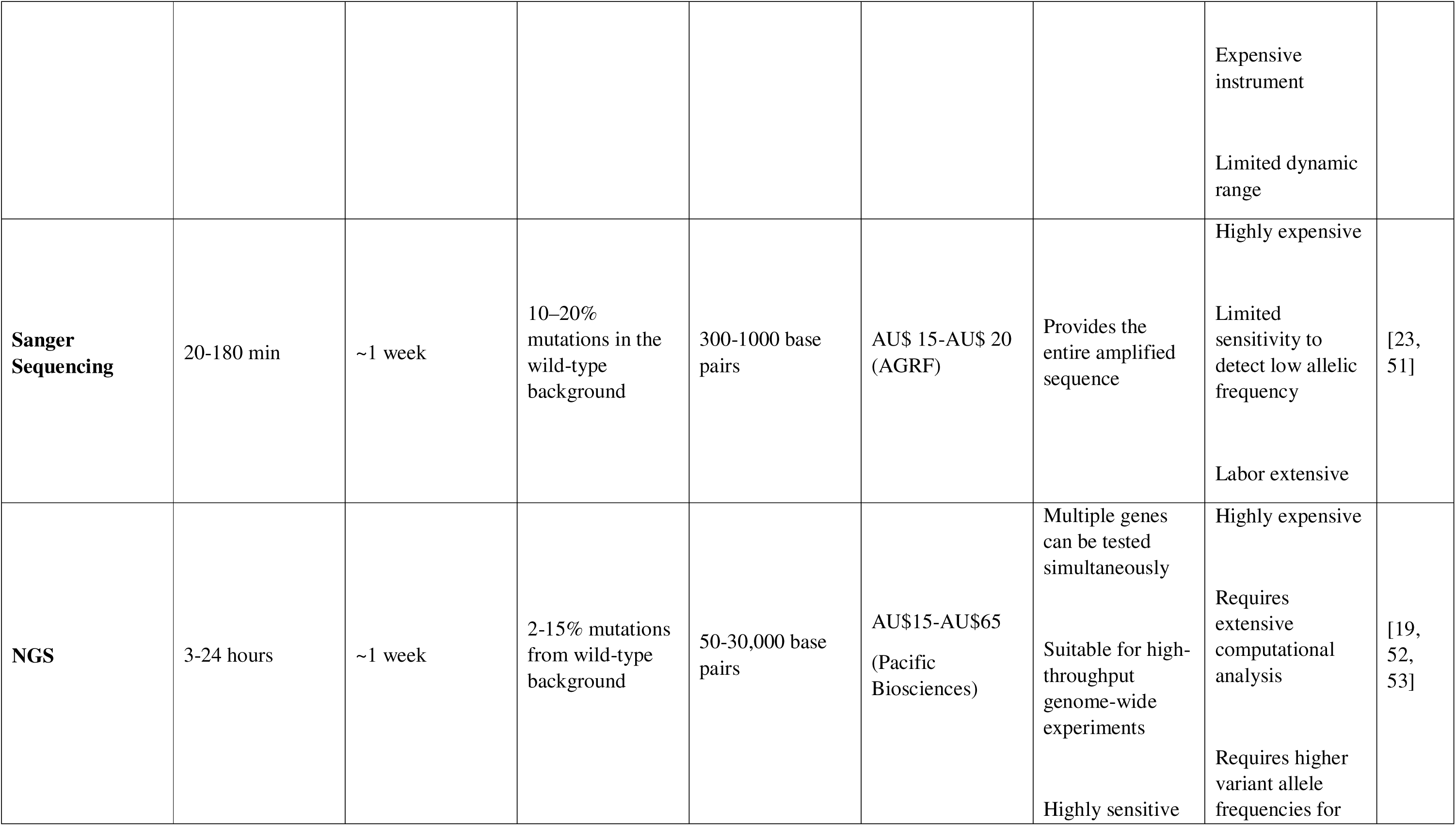
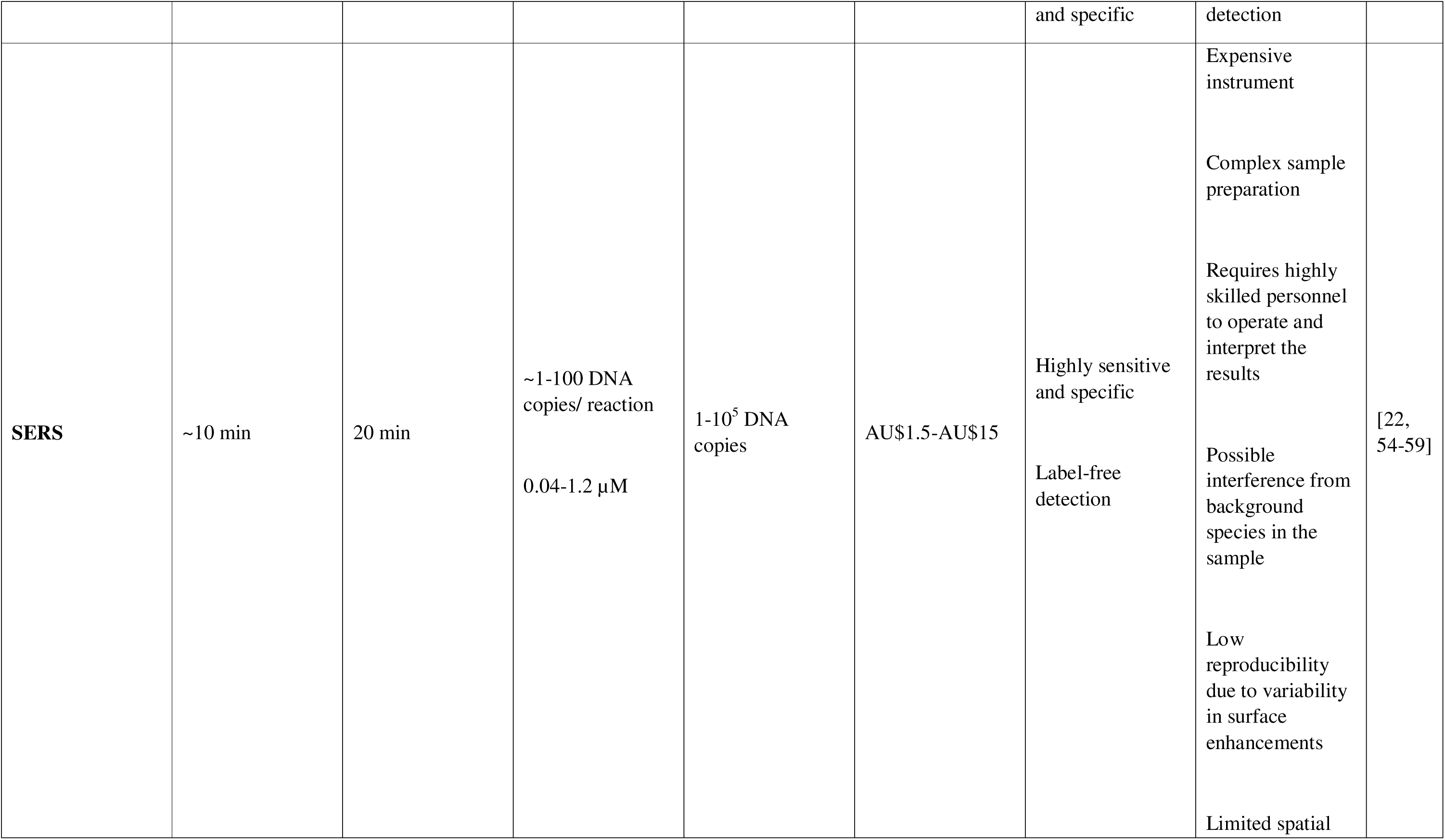
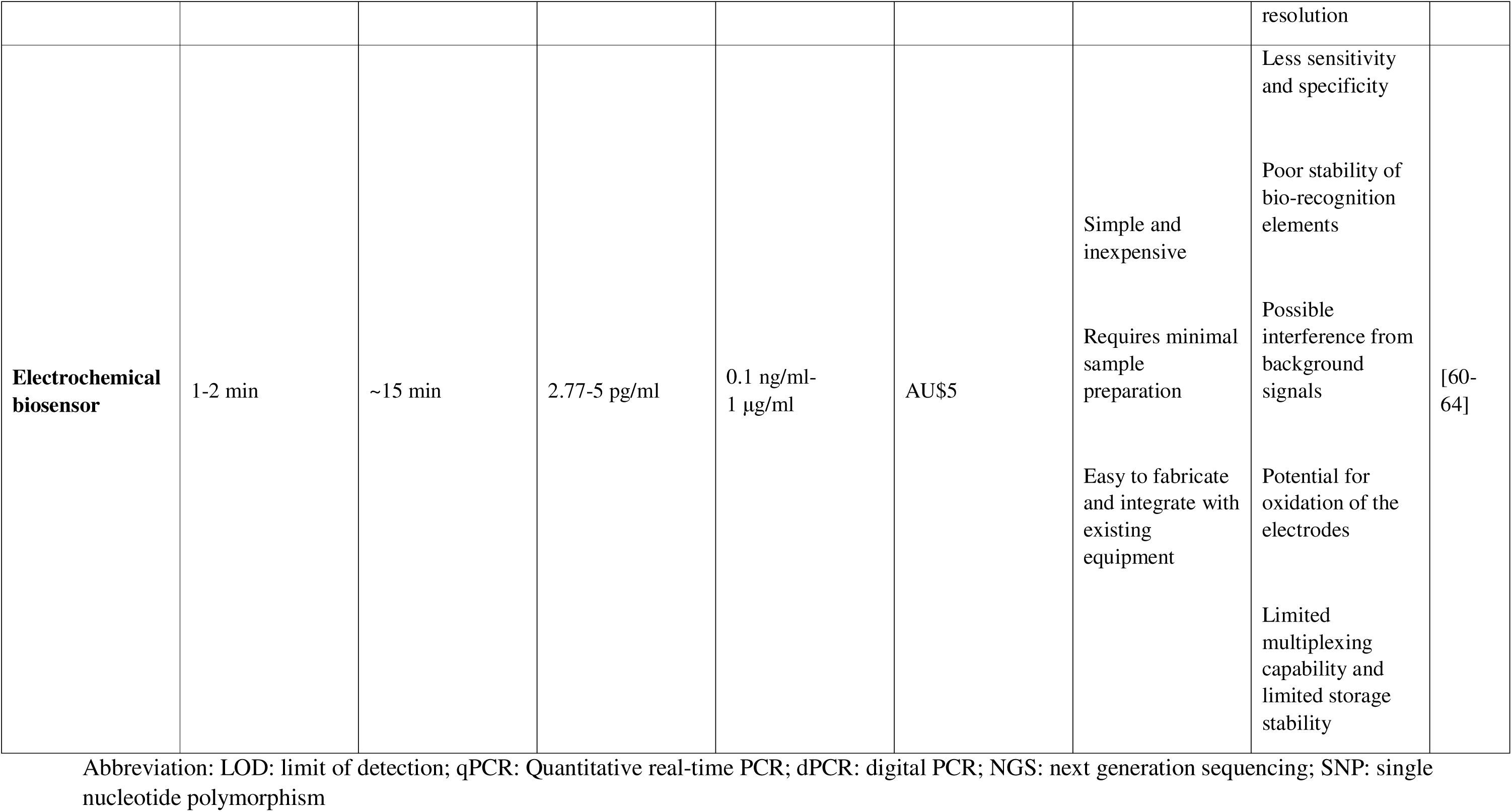
Comparison of analytical performance between PNA-LNA molecular switch and contemporary standard mutation detection methods.

Some discordant results were observed in our analysis. Specifically, in two cases from each panel (*KRAS* and *BRAF*), mutations were detected in the CTCs despite the corresponding tissue samples being wild-type. Conversely, six cases in the *KRAS* panel and one case in the *BRAF* panel exhibited mutations in the primary tumors that were not detected in the CTCs. For the remaining samples, a significant correlation was observed (*p*<0.001). These discrepancies may be explained by the presence of genetically distinct CTC subclones, arising from intratumor heterogeneity, that enter the bloodstream and differ from the primary tumor. Such subclones may compete during the shedding of tumour cells into circulation, a phenomenon supported by previous studies [13, 30, 31]. Another possible explanation lies in the differing detection thresholds of the two methods for identifying mutations based on Variant Allele Frequency (VAF). However, due to the unavailability of VAF data in this study, this could not be confirmed. Additionally, during tumour progression and evolution, CTCs may acquire new mutations, leading to genotypic differences from their matched primary tumors. Lastly, due to resource limitations, our analysis was based on heterogeneous CTC populations, which may have included some leukocyte contamination. This could have introduced confounding effects and impacted the accuracy of mutation detection to some extent.

There was no significant concordance between the *KRAS* oncogene mutation status with the different clinical pathological parameters in the primary adenocarcinoma. However, a notable trend was observed: most mutations were identified in tumors with advanced local spread (19 of 26 cases) and in higher pathological stages (17 of 26 cases). Additionally, CRCs with high MSI exhibited fewer *KRAS* mutations (4 of 26 cases). These findings are consistent with previous research, which suggests that *KRAS* mutations promote aggressive tumor progression[32-34]. Similarly, *BRAF* mutations were more frequently observed in tumors with advanced local spread (15 of 19 cases) and were less common in high MSI phenotypes (7 of 19 cases). Despite these observations, no significant concordance was found between *BRAF* mutations and clinical parameters. Consistent with earlier studies, *KRAS* mutations were more prevalent than *BRAF* mutations among the recruited adenocarcinomas. Furthermore, all but one of the samples exhibited mutually exclusive mutations in *KRAS* or *BRAF*, reflecting the well-documented mutual exclusivity of these mutations in CRC[35-37].

Unlike the primary adenocarcinoma, CTCs exhibited significant correlations between *KRAS* mutations and certain clinicopathological parameters, including lymph node metastasis, overall pathological stage, and LVI. Out of 13 CTC samples that harboured mutations in the *KRAS* gene, 11 cases had lymph node metastasis (*p*=0.002), which is a well-established prognostic factor for predicting disease recurrence and survival in CRC patients[38]. Since tumor cells are thought to spread from the primary site to distant organs via lymph nodes and lymphatic vessels, local lymph node metastasis is considered a critical step in tumor cell dissemination in CRC. Similarly, LVI is another important indicator of the aggressiveness of CRC, reflecting the tumour’s interaction with lymphatic or vascular channels—an essential prerequisite for metastasis [39, 40]. In our study, *KRAS* mutations were more prevalent in patients with positive LVI (9 vs 4), showing a significant correlation (*p*=0.034). These findings underscore the potential role of *KRAS* mutations in promoting metastatic behaviour through LVI and lymph node involvement in CRC. Furthermore, the overall pathological stage (TNM staging) helps in identifying the anatomic extent of cancer and serves as a measure of disease severity, with advanced staging being associated with decreased survivability[41]. We found that 85% of the mutations (11 vs 2) in the *KRAS* gene from CTCs were from patients with advanced pathological stages (*p*=0.005). These results support the hypothesis that CTCs with *KRAS* mutations may originate from advanced and aggressive tumors. However, like the adenocarcinoma cases, none of the *BRAF* mutations in CTCs correlated significantly with the clinical characteristics of the recruited CRC patients. This lack of correlation may be attributed to the smaller number of CTC samples and the lower frequency of *BRAF* mutations in the cohort studied.

To validate the results of the PNA-LNA molecular switch, we compared them with NGS data from tissue samples of 27 patients for both *KRAS* and *BRAF* genes. We observed highly concordant results between the two methods, with only three cases in each panel yielding conflicting outcomes (*p*<0.001). These discrepancies may be due to the differing sizes of the tumor blocks used in each method, which may have contained distinct genetic subclones of the tumor. Despite the limitation of the PNA-LNA molecular switch in detecting only known SNPs, the significant concordance with NGS supports its potential for clinical use. Given its sensitivity, comparability to traditional detection techniques, and rapid, cost-effective nature, this method could be suitable for point-of-care applications in the future.

While the method shows considerable promise, it is not without limitations. Its current design restricts analysis to known SNPs, limiting its ability to detect novel or uncharacterized mutations. Each reaction targets a single genetic region, which can reduce efficiency when multiple mutations need to be assessed. The approach also relies on commercially synthesized PNA and LNA probes, contributing to higher costs and increased dependence on external suppliers. Additionally, the method is less suitable for analyzing circulating tumor DNA (ctDNA) and cell-free DNA (cfDNA), which are often highly fragmented and may hinder accurate mutation detection. To address these challenges, ongoing work is focused on enhancing the method’s multiplexing capacity and improving its compatibility with fragmented DNA. In parallel, validation studies with larger patient cohorts are underway to further establish its clinical utility and support broader implementation. Looking ahead, efforts are also being directed toward integrating this approach into a portable, point-of-care diagnostic device, which could offer rapid and accessible mutation detection in diverse clinical settings.

## Conclusion

In summary, we employed the PNA-LNA molecular switch to detect the mutation status of *KRAS* (G12V) and *BRAF* (V600E) genes in primary adenocarcinoma and CTCs from patients with CRC. Our analysis revealed significant correlations between *KRAS* mutations in CTCs and clinical parameters such as lymph node metastasis, overall pathological stage, and LVI. Furthermore, the results from the PNA-LNA molecular switch showed significant concordance with the NGS data, supporting the feasibility of this method for clinical applications.

## CRediT Author Contribution

**M.S.I:** Conceptualization, Methodology, Software, Validation, Formal analysis, Investigation, Data curation, Writing-Original Draft, Visualization, Project administration. **E.R:** Formal analysis, Investigation, Data curation. **S.A:** Investigation, Data curation. **N.M:** Investigation, Data curation. **C.T.L:** Resources. **M.J.A.S:** Conceptualization, Methodology, Writing - Review & Editing, Supervision. **V.G:** Conceptualization, Methodology, Writing - Review & Editing, Supervision, Project administration. **A.K.L:** Resources, Writing - Review & Editing, Supervision, Project administration.

## Data availability statement

Data will be available upon request.

## Ethics approval statement

Ethical approval was obtained from the Griffith University Human Research Ethics Committee (GU Ref No: MSC/17/10/HREC).

## Patient consent statement

Written informed consent was obtained from each patient.

## Acknowledgment

M.S.I. acknowledges the support from Griffith University Postgraduate Research Scholarship and conveys his gratitude to **Tanzena Tanny** for sharing Biorender.

## References

[1] S. Alinia, S. Ahmadi, Z. Mohammadi, F. Rastkar Shirvandeh, M. Asghari-Jafarabadi, L. Mahmoudi, M. Safari, G. Roshanaei, Exploring the impact of stage and tumor site on colorectal cancer survival: Bayesian survival modeling, Sci. Rep., 14 (2024) 4270.

[2] Colorectal Cancer Early Detection, Diagnosis, and Staging, in, vol. 2024, American Cancer Society, 2022.

[3] C. Therkildsen, T.K. Bergmann, T. Henrichsen-Schnack, S. Ladelund, M. Nilbert, The predictive value of KRAS, NRAS, BRAF, PIK3CA and PTEN for anti-EGFR treatment in metastatic colorectal cancer: A systematic review and meta-analysis, Acta Oncol, 53 (2014) 852–864.

[4] X.-J. Luo, Q. Zhao, J. Liu, J.-B. Zheng, M.-Z. Qiu, H.-Q. Ju, R.-H. Xu, Novel Genetic and Epigenetic Biomarkers of Prognostic and Predictive Significance in Stage II/III Colorectal Cancer, Mol. Ther., 29 (2021) 587–596.

[5] M. Röring, T. Brummer, Aberrant B-Raf signaling in human cancer -- 10 years from bench to bedside, Crit Rev Oncog, 17 (2012) 97–121.

[6] E. Oikonomou, E. Koustas, M. Goulielmaki, A. Pintzas, BRAF vs RAS oncogenes: Are mutations of the same pathway equal? Differential signalling and therapeutic implications, Oncotarget, 5 (2014).

[7] M.J. Sorich, M.D. Wiese, A. Rowland, G. Kichenadasse, R.A. McKinnon, C.S. Karapetis, Extended RAS mutations and anti-EGFR monoclonal antibody survival benefit in metastatic colorectal cancer: a meta-analysis of randomized, controlled trials, Ann. Oncol., 26 (2015) 13–21.

[8] E.M.J. van Brummelen, A. de Boer, J.H. Beijnen, J.H.M. Schellens, BRAF Mutations as Predictive Biomarker for Response to Anti[EGFR Monoclonal Antibodies, Oncologist, 22 (2017) 864–872.

[9] K.M. Haigis, KRAS Alleles: The Devil Is in the Detail, Trends Cancer, 3 (2017) 686–697.

[10] E. Sanz-Garcia, G. Argiles, E. Elez, J. Tabernero, BRAF mutant colorectal cancer: prognosis, treatment, and new perspectives, Ann. Oncol., 28 (2017) 2648–2657.

[11] P. Liu, Y. Wang, X. Li, Targeting the untargetable KRAS in cancer therapy, Acta Pharm Sin B, 9 (2019) 871–879.

[12] L. Ung, A.K. Lam, D.L. Morris, T.C. Chua, Tissue-based biomarkers predicting outcomes in metastatic colorectal cancer: a review, Clin Transl Oncol, 16 (2014) 425–435.

[13] Q. Wang, L. Zhao, L. Han, X. Tuo, S. Ma, Y. Wang, X. Feng, D. Liang, C. Sun, Q. Wang, Q. Song, Q. Li, The Discordance of Gene Mutations between Circulating Tumor Cells and Primary/Metastatic Tumor, Mol Ther Oncolytics, 15 (2019) 21–29.

[14] D.H. Baek, G.H. Kim, G.A. Song, I.S. Han, E.Y. Park, H.S. Kim, H.J. Jo, S.H. Ko, D.Y. Park, Y.K. Cho, Clinical Potential of Circulating Tumor Cells in Colorectal Cancer: A Prospective Study, Clin Transl Gastroenterol, 10 (2019) e00055.

[15] A. Kalikaki, H. Politaki, J. Souglakos, S. Apostolaki, E. Papadimitraki, N. Georgoulia, M. Tzardi, D. Mavroudis, V. Georgoulias, A. Voutsina, KRAS Genotypic Changes of Circulating Tumor Cells during Treatment of Patients with Metastatic Colorectal Cancer, PLoS One, 9 (2014) e104902.

[16] A.K. Casasent, A. Schalck, R. Gao, E. Sei, A. Long, W. Pangburn, T. Casasent, F. Meric-Bernstam, M.E. Edgerton, N.E. Navin, Multiclonal Invasion in Breast Tumors Identified by Topographic Single Cell Sequencing, Cell, 172 (2018) 205–217.e212.

[17] J. Zhang, J. Fujimoto, J. Zhang, D.C. Wedge, X. Song, J. Zhang, S. Seth, C.-W. Chow, Y. Cao, C. Gumbs, K.A. Gold, N. Kalhor, L. Little, H. Mahadeshwar, C. Moran, A. Protopopov, H. Sun, J. Tang, X. Wu, Y. Ye, W.N. William, J.J. Lee, J.V. Heymach, W.K. Hong, S. Swisher, I.I. Wistuba, P.A. Futreal, Intratumor heterogeneity in localized lung adenocarcinomas delineated by multiregion sequencing, Science, 346 (2014) 256–259.

[18] J.-J. Hao, D.-C. Lin, H.Q. Dinh, A. Mayakonda, Y.-Y. Jiang, C. Chang, Y. Jiang, C.-C. Lu, Z.-Z. Shi, X. Xu, Y. Zhang, Y. Cai, J.-W. Wang, Q.-M. Zhan, W.-Q. Wei, B.P. Berman, M.-R. Wang, H.P. Koeffler, Spatial intratumoral heterogeneity and temporal clonal evolution in esophageal squamous cell carcinoma, Nat. Genet., 48 (2016) 1500–1507.

[19] J. Chung, S. Xiao, Y. Gao, Y.H. Soung, Recent Technologies towards Diagnostic and Therapeutic Applications of Circulating Nucleic Acids in Colorectal Cancers, Int J Mol Sci, 25 (2024) 8703.

[20] M.S. Islam, S. Aktar, N. Moetamedirad, N. Xie, C.T. Lu, V. Gopalan, A.K. Lam, M.J.A. Shiddiky, A novel platform for mutation detection in colorectal cancer using a PNA-LNA molecular switch, Biosensors Bioelectron., 267 (2025) 116813.

[21] J. Li, Z. Gao, J. Chen, R. Cheng, J. Niu, J. Zhang, Y. Yang, X. Yuan, J. Xia, G. Mao, H. Liu, Y. Dong, C. Wu, Development of a panel of three multiplex allele-specific qRT-PCR assays for quick differentiation of recombinant variants and Omicron subvariants of SARS-CoV-2, Front Cell Infect Microbiol, 12 (2022) 953027.

[22] Y. Liu, N. Lyu, V.K. Rajendran, J. Piper, A. Rodger, Y. Wang, Sensitive and Direct DNA Mutation Detection by Surface-Enhanced Raman Spectroscopy Using Rational Designed and Tunable Plasmonic Nanostructures, Analytical Chemistry, 92 (2020) 5708–5716.

[23] R.E. Shackelford, N.A. Whitling, P. McNab, S. Japa, D. Coppola, KRAS Testing:A Tool for the Implementation of Personalized Medicine, Genes Cancer, 3 (2012) 459–466.

[24] X. Sun, K. Hung, L. Wu, D. Sidransky, B. Guo, Detection of tumor mutations in the presence of excess amounts of normal DNA, Nat Biotechnol, 20 (2002) 186–189.

[25] S. Obika, D. Nanbu, Y. Hari, K.-i. Morio, Y. In, T. Ishida, T. Imanishi, Synthesis of 2′-O,4′-C-methyleneuridine and -cytidine. Novel bicyclic nucleosides having a fixed C3, -endo sugar puckering, Tetrahedron Lett., 38 (1997) 8735–8738.

[26] I. Nagtegaal, M. Arends, M. Salto-Tellez, Colorectal adenocarcinoma, in: I. Nagtegaal, M. Arends, R. Odze, A. Lam (Eds.) Tumours of the colon and rectum, International Agency for Research on Cancer, 2019, pp. 177–187.

[27] S. Aktar, S.M.K. Gamage, T. Cheng, N. Parkneshan, C.T. Lu, F. Islam, V. Gopalan, A.K.-y. Lam, Gene Expression Analysis of Immune Regulatory Genes in Circulating Tumour Cells and Peripheral Blood Mononuclear Cells in Patients with Colorectal Carcinoma, Int J Mol Sci, 24 (2023) 5051.

[28] F.B. Hamid, C.T. Lu, M. Matos, T. Cheng, V. Gopalan, A.K. Lam, Enumeration, characterisation and clinicopathological significance of circulating tumour cells in patients with colorectal carcinoma, Cancer Genet, 254–255 (2021) 48–57.

[29] S. Aktar, F. Islam, T. Cheng, S.M.K. Gamage, I.N. Choudhury, M.S. Islam, C.T. Lu, F.B. Hamid, H. Ishida, I. Abe, N. Xie, V. Gopalan, A.K. Lam, Correlation between KRAS Mutation and CTLA-4 mRNA Expression in Circulating Tumour Cells: Clinical Implications in Colorectal Cancer, Genes, 14 (2023) 1808.

[30] C. Raimondi, C. Nicolazzo, A. Gradilone, G. Giannini, E. De Falco, I. Chimenti, E. Varriale, S. Hauch, L. Plappert, E. Cortesi, P. Gazzaniga, Circulating tumor cells, Cancer Biol Ther, 15 (2014) 496–503.

[31] C. Zhang, Y. Guan, Y. Sun, D. Ai, Q. Guo, Tumor heterogeneity and circulating tumor cells, Cancer Lett., 374 (2016) 216–223.

[32] F. El Agy, S. el Bardai, I. El Otmani, Z. Benbrahim, I.M.H. Karim, K. Mazaz, E.B. Benjelloun, A. Ousadden, M. El Abkari, S.A. Ibrahimi, L. Chbani, Mutation status and prognostic value of KRAS and NRAS mutations in Moroccan colon cancer patients: A first report, PLoS One, 16 (2021) e0248522.

[33] A. Mannan, V. Hahn-Strömberg, K-ras mutations are correlated to lymph node metastasis and tumor stage, but not to the growth pattern of colon carcinoma, APMIS, 120 (2012) 459–468.

[34] J. Rimbert, G. Tachon, A. Junca, C. Villalva, L. Karayan-Tapon, D. Tougeron, Association between clinicopathological characteristics and RAS mutation in colorectal cancer, Mod Pathol, 31 (2018) 517–526.

[35] D.M. Muzny, M.N. Bainbridge, K. Chang, H.H. Dinh, J.A. Drummond, G. Fowler, C.L. Kovar, L.R. Lewis, M.B. Morgan, I.F. Newsham, J.G. Reid, J. Santibanez, E. Shinbrot, L.R. Trevino, Y.-Q. Wu, M. Wang, P. Gunaratne, L.A. Donehower, C.J. Creighton, D.A. Wheeler, R.A. Gibbs, M.S. Lawrence, D. Voet, R. Jing, K. Cibulskis, A. Sivachenko, P. Stojanov, A. McKenna, E.S. Lander, S. Gabriel, G. Getz, L. Ding, R.S. Fulton, D.C. Koboldt, T. Wylie, J. Walker, D.J. Dooling, L. Fulton, K.D. Delehaunty, C.C. Fronick, R. Demeter, E.R. Mardis, R.K. Wilson, A. Chu, H.-J.E. Chun, A.J. Mungall, E. Pleasance, A. Gordon Robertson, D. Stoll, M. Balasundaram, I. Birol, Y.S.N. Butterfield, E. Chuah, R.J.N. Coope, N. Dhalla, R. Guin, C. Hirst, M. Hirst, R.A. Holt, D. Lee, H.I. Li, M. Mayo, R.A. Moore, J.E. Schein, J.R. Slobodan, A. Tam, N. Thiessen, R. Varhol, T. Zeng, Y. Zhao, S.J.M. Jones, M.A. Marra, A.J. Bass, A.H. Ramos, G. Saksena, A.D. Cherniack, S.E. Schumacher, B. Tabak, S.L. Carter, N.H. Pho, H. Nguyen, R.C. Onofrio, A. Crenshaw, K. Ardlie, R. Beroukhim, W. Winckler, G. Getz, M. Meyerson, A. Protopopov, J. Zhang, A. Hadjipanayis, E. Lee, R. Xi, L. Yang, X. Ren, H. Zhang, N. Sathiamoorthy, S. Shukla, P.-C. Chen, P. Haseley, Y. Xiao, S. Lee, J. Seidman, L. Chin, P.J. Park, R. Kucherlapati, J. Todd Auman, K.A. Hoadley, Y. Du, M.D. Wilkerson, Y. Shi, C. Liquori, S. Meng, L. Li, Y.J. Turman, M.D. Topal, D. Tan, S. Waring, E. Buda, J. Walsh, C.D. Jones, P.A. Mieczkowski, D. Singh, J. Wu, A. Gulabani, P. Dolina, T. Bodenheimer, A.P. Hoyle, J.V. Simons, M. Soloway, L.E. Mose, S.R. Jefferys, S. Balu, B.D. O’Connor, J.F. Prins, D.Y. Chiang, D. Neil Hayes, C.M. Perou, T. Hinoue, D.J. Weisenberger, D.T. Maglinte, F. Pan, B.P. Berman, D.J. Van Den Berg, H. Shen, T. Triche Jr, S.B. Baylin, P.W. Laird, G. Getz, M. Noble, D. Voet, G. Saksena, N. Gehlenborg, D. DiCara, J. Zhang, H. Zhang, C.-J. Wu, S. Yingchun Liu, S. Shukla, M.S. Lawrence, L. Zhou, A. Sivachenko, P. Lin, P. Stojanov, R. Jing, R.W. Park, M.-D. Nazaire, J. Robinson, H. Thorvaldsdottir, J. Mesirov, P.J. Park, L. Chin, V. Thorsson, S.M. Reynolds, B. Bernard, R. Kreisberg, J. Lin, L. Iype, R. Bressler, T. Erkkilä, M. Gundapuneni, Y. Liu, A. Norberg, T. Robinson, D. Yang, W. Zhang, I. Shmulevich, J.J. de Ronde, N. Schultz, E. Cerami, G. Ciriello, A.P. Goldberg, B. Gross, A. Jacobsen, J. Gao, B. Kaczkowski, R. Sinha, B. Arman Aksoy, Y. Antipin, B. Reva, R. Shen, B.S. Taylor, T.A. Chan, M. Ladanyi, C. Sander, R. Akbani, N. Zhang, B.M. Broom, T. Casasent, A. Unruh, C. Wakefield, S.R. Hamilton, R. Craig Cason, K.A. Baggerly, J.N. Weinstein, D. Haussler, C.C. Benz, J.M. Stuart, S.C. Benz, J. Zachary Sanborn, C.J. Vaske, J. Zhu, C. Szeto, G.K. Scott, C. Yau, S. Ng, T. Goldstein, K. Ellrott, E. Collisson, A.E. Cozen, D. Zerbino, C. Wilks, B. Craft, P. Spellman, R. Penny, T. Shelton, M. Hatfield, S. Morris, P. Yena, C. Shelton, M. Sherman, J. Paulauskis, J.M. Gastier-Foster, J. Bowen, N.C. Ramirez, A. Black, R. Pyatt, L. Wise, P. White, M. Bertagnolli, J. Brown, T.A. Chan, G.C. Chu, C. Czerwinski, F. Denstman, R. Dhir, A. Dörner, C.S. Fuchs, J.G. Guillem, M. Iacocca, H. Juhl, A. Kaufman, B. Kohl Iii, X. Van Le, M.C. Mariano, E.N. Medina, M. Meyers, G.M. Nash, P.B. Paty, N. Petrelli, B. Rabeno, W.G. Richards, D. Solit, P. Swanson, L. Temple, J.E. Tepper, R. Thorp, E. Vakiani, M.R. Weiser, J.E. Willis, G. Witkin, Z. Zeng, M.J. Zinner, N. The Cancer Genome Atlas, M. Genome Sequencing Center Baylor College of, I. Genome Sequencing Center Broad, L. Genome Sequencing Center Washington University in St, B.C.C.A. Genome Characterization Center, I. Genome-Characterization Center Broad, B. Genome-Characterization Center, H. Women’s, S. Harvard Medical, C.H. Genome-Characterization Center University of North Carolina, C. Genome-Characterization Centers University of Southern, U. Johns Hopkins, I. Genome Data Analysis Center Broad, B. Genome Data Analysis Center Institute for Systems, C. Genome Data Analysis Center Memorial Sloan-Kettering Cancer, M.D.A.C.C. Genome Data Analysis Center University of Texas, U.o.C.S.C. Genome Data Analysis Centers, I. the Buck, C. Biospecimen Core Resource International Genomics, R. Nationwide Children’s Hospital Biospecimen Core, s. Tissue source, g. disease working, Comprehensive molecular characterization of human colon and rectal cancer, Nature, 487 (2012) 330–337.

[36] S.A. Forbes, D. Beare, P. Gunasekaran, K. Leung, N. Bindal, H. Boutselakis, M. Ding, S. Bamford, C. Cole, S. Ward, C.Y. Kok, M. Jia, T. De, J.W. Teague, M.R. Stratton, U. McDermott, P.J. Campbell, COSMIC: exploring the world’s knowledge of somatic mutations in human cancer, Nucleic Acids Res., 43 (2014) D805–D811.

[37] M. Morkel, P. Riemer, H. Bläker, C. Sers, Similar but different: distinct roles for KRAS and BRAF oncogenes in colorectal cancer development and therapy resistance, Oncotarget, 6 (2015) 20785–20800.

[38] H.J. Kim, G.S. Choi, Clinical Implications of Lymph Node Metastasis in Colorectal Cancer: Current Status and Future Perspectives, Ann Coloproctol, 35 (2019) 109–117.

[39] L. Zhang, Y. Deng, S. Liu, W. Zhang, Z. Hong, Z. Lu, Z. Pan, X. Wu, J. Peng, Lymphovascular invasion represents a superior prognostic and predictive pathological factor of the duration of adjuvant chemotherapy for stage III colon cancer patients, BMC Cancer, 23 (2023) 3.

[40] Y. Akagi, Y. Adachi, T. Ohchi, T. Kinugasa, K. Shirouzu, Prognostic Impact of Lymphatic Invasion of Colorectal Cancer: A Single-center Analysis of 1,616 Patients Over 24 Years, Anticancer Res., 33 (2013) 2965.

[41] G. Menon, A. Recio-Boiles, S. Lotfollahzadeh, B. Cagir, Colon Cancer, in: StatPearls, StatPearls Publishing, Treasure Island (FL), 2024.

[42] N. Soda, B.H.A. Rehm, P. Sonar, N.-T. Nguyen, M.J.A. Shiddiky, Advanced liquid biopsy technologies for circulating biomarker detection, Journal of Materials Chemistry B, 7 (2019) 6670–6704.

[43] Z. Xue, M. You, P. Peng, H. Tong, W. He, A. Li, P. Mao, T. Xu, F. Xu, C. Yao, Taqman-MGB nanoPCR for Highly Specific Detection of Single-Base Mutations, Int J Nanomedicine, 16 (2021) 3695–3705.

[44] A.C. McEvoy, B.A. Wood, N.M. Ardakani, M.R. Pereira, R. Pearce, L. Cowell, C. Robinson, F. Grieu-Iacopetta, A.J. Spicer, B. Amanuel, M. Ziman, E.S. Gray, Droplet Digital PCR for Mutation Detection in Formalin-Fixed, Paraffin-Embedded Melanoma Tissues: A Comparison with Sanger Sequencing and Pyrosequencing, The Journal of Molecular Diagnostics, 20 (2018) 240–252.

[45] A.F. Al Mana, K. Culp, A. Keeler, O. Perrera, M. Rajagopalan, L. Jacky, B. Brown, B. Thyagarajan, Performance of a Rapid Digital PCR Test for the Detection of Non-Small Cell Lung Cancer (NSCLC) Variants, Molecular Diagnosis & Therapy, (2024).

[46] S. Galbiati, F. Damin, L. Ferraro, N. Soriani, V. Burgio, M. Ronzoni, L. Gianni, M. Ferrari, M. Chiari, Microarray Approach Combined with ddPCR: An Useful Pipeline for the Detection and Quantification of Circulating Tumour dna Mutations, Cells, 8 (2019).

[47] C.A. Milbury, Q. Zhong, J. Lin, M. Williams, J. Olson, D.R. Link, B. Hutchison, Determining lower limits of detection of digital PCR assays for cancer-related gene mutations, Biomolecular Detection and Quantification, 1 (2014) 8–22.

[48] X. Luo, K. Wang, Y. Xue, X. Cao, J. Zhou, J. Wang, Digital PCR-free technologies for absolute quantitation of nucleic acids at single-molecule level, Chinese Chemical Letters, (2024) 109924.

[49] J. Kuypers, K.R. Jerome, Applications of Digital PCR for Clinical Microbiology, Journal of Clinical Microbiology, 55 (2017) 1621–1628.

[50] M.H. Cleveland, H.-J. He, M. Milavec, Y.-K. Bae, P.M. Vallone, J.F. Huggett, Digital PCR for the characterization of reference materials, Molecular Aspects of Medicine, 96 (2024) 101256.

[51] Y.H. Yan, S.X. Chen, L.Y. Cheng, A.Y. Rodriguez, R. Tang, K. Cabrera, D.Y. Zhang, Confirming putative variants at[≤[5% allele frequency using allele enrichment and Sanger sequencing, Sci. Rep., 11 (2021) 11640.

[52] K. Turabi, K. Klute, P. Radhakrishnan, Decoding the Dynamics of Circulating Tumor DNA in Liquid Biopsies, Cancers, 16 (2024) 2432.

[53] T. Miura, S. Yasuda, Y. Sato, A simple method to estimate the in-house limit of detection for genetic mutations with low allele frequencies in whole-exome sequencing analysis by next-generation sequencing, BMC Genomic Data, 22 (2021) 8.

[54] Y. Liu, N. Lyu, V.K. Rajendran, J. Piper, A. Rodger, Y. Wang, Sensitive and Direct DNA Mutation Detection by Surface-Enhanced Raman Spectroscopy Using Rational Designed and Tunable Plasmonic Nanostructures, Anal Chem, 92 (2020) 5708–5716.

[55] A.V. Markin, A.I. Arzhanukhina, N.E. Markina, I.Y. Goryacheva, Analytical performance of electrochemical surface-enhanced Raman spectroscopy: A critical review, TrAC Trends in Analytical Chemistry, 157 (2022) 116776.

[56] S. Sloan-Dennison, E. O’Connor, J.W. Dear, D. Graham, K. Faulds, Towards quantitative point of care detection using SERS lateral flow immunoassays, Analytical and Bioanalytical Chemistry, 414 (2022) 4541–4549.

[57] S. Yan, F. Chu, H. Zhang, Y. Yuan, Y. Huang, A. Liu, S. Wang, W. Li, S. Li, W. Wen, Rapid, one-step preparation of SERS substrate in microfluidic channel for detection of molecules and heavy metal ions, Spectrochimica Acta Part A: Molecular and Biomolecular Spectroscopy, 220 (2019) 117113.

[58] H. Marks, M. Schechinger, J. Garza, A. Locke, G. Coté, Surface enhanced Raman spectroscopy (SERS) for in vitro diagnostic testing at the point of care, Nanophotonics, 6 (2017) 681–701.

[59] J. Langer, D. Jimenez de Aberasturi, J. Aizpurua, R.A. Alvarez-Puebla, B. Auguié, J.J. Baumberg, G.C. Bazan, S.E.J. Bell, A. Boisen, A.G. Brolo, J. Choo, D. Cialla-May, V. Deckert, L. Fabris, K. Faulds, F.J. García de Abajo, R. Goodacre, D. Graham, A.J. Haes, C.L. Haynes, C. Huck, T. Itoh, M. Käll, J. Kneipp, N.A. Kotov, H. Kuang, E.C. Le Ru, H.K. Lee, J.-F. Li, X.Y. Ling, S.A. Maier, T. Mayerhöfer, M. Moskovits, K. Murakoshi, J.-M. Nam, S. Nie, Y. Ozaki, I. Pastoriza-Santos, J. Perez-Juste, J. Popp, A. Pucci, S. Reich, B. Ren, G.C. Schatz, T. Shegai, S. Schlücker, L.-L. Tay, K.G. Thomas, Z.-Q. Tian, R.P. Van Duyne, T. Vo-Dinh, Y. Wang, K.A. Willets, C. Xu, H. Xu, Y. Xu, Y.S. Yamamoto, B. Zhao, L.M. Liz-Marzán, Present and Future of Surface-Enhanced Raman Scattering, ACS Nano, 14 (2020) 28–117.

[60] W. Lu, S. Xue, X. Liu, C. Bao, H. Shi, An ultrasensitive photoelectrochemical immunosensor for carcinoembryonic antigen detection based on cobalt borate nanosheet array and Mn2O3@Au nanocube, Microchemical Journal, 196 (2024) 109606.

[61] M.R. Hasan, M.S. Ahommed, M. Daizy, M.S. Bacchu, M.R. Ali, M.R. Al-Mamun, M.A. Saad Aly, M.Z.H. Khan, S.I. Hossain, Recent development in electrochemical biosensors for cancer biomarkers detection, Biosensors and Bioelectronics: X, 8 (2021) 100075.

[62] S.N. Topkaya, M. Azimzadeh, M. Ozsoz, Electrochemical Biosensors for Cancer Biomarkers Detection: Recent Advances and Challenges, Electroanalysis, 28 (2016) 1402–1419.

[63] S. Menon, M.R. Mathew, S. Sam, K. Keerthi, K.G. Kumar, Recent advances and challenges in electrochemical biosensors for emerging and re-emerging infectious diseases, J Electroanal Chem (Lausanne), 878 (2020) 114596.

[64] C. Bao, R. Zhang, Y. Qiao, X. Cao, F. He, W. Hu, M. Wei, W. Lu, Au Nanoparticles Anchored on Cobalt Boride Nanowire Arrays for the Electrochemical Determination of Prostate-Specific Antigen, ACS Applied Nano Materials, 4 (2021) 5707–5716.

