## Supplementary Table S1 for "Mutation profiling of *KRAS* and *BRAF* in primary tumors and circulating tumor cells of colorectal cancer patients using PNA-LNA molecular switch"

**Supplementary Table S1:** Mutation status of patients for *KRAS* (G12V) and *BRAF* (V600E) from primary adenocarcinoma and matched CTCs with PNA-LNA molecular switch and NGS

| **Patient no.** | **KRAS Tissue (G12V) LAMP** | **BRAF Tissue (V600E) LAMP** | **#CTC** | **KRAS CTC (G12V) LAMP** | **BRAF CTC (V600E) LAMP** | **KRAS Sequencing** | **BRAF Sequencing** |
| --- | --- | --- | --- | --- | --- | --- | --- |
| Patient 1 | 0 | 1 | - | - | - | - | - |
| Patient 2 | 1 | 0 | - | - | - | - | - |
| Patient 3 | 0 | 1 | - | - | - | - | - |
| Patient 4 | 0 | 0 | - | - | - | 1 | 0 |
| Patient 5 | 0 | 1 | - | - | - | - | - |
| Patient 6 | 0 | 0 | - | - | - | 0 | 0 |
| Patient 7 | 0 | 1 | - | - | - | - | - |
| Patient 8 | 0 | 0 | - | - | - | - | - |
| Patient 9 | 0 | 0 | - | - | - | - | - |
| Patient 10 | 1 | 0 | - | - | - | 1 | 0 |
| Patient 11 | 0 | 1 | - | - | - | - | - |
| Patient 12 | 0 | 0 | - | - | - | - | - |
| Patient 13 | 0 | 1 | - | - | - | - | - |
| Patient 14 | 0 | 1 | - | - | - | 0 | 0 |
| Patient 15 | 1 | 1 | 32 | 0 | 0 | - | - |
| Patient 16 | 0 | 0 | 12 | 1 | 0 | - | - |
| Patient 17 | 0 | 0 | 14 | 0 | 0 | 0 | 0 |
| Patient 18 | 0 | 0 | 4 | 0 | 1 | 0 | 0 |
| Patient 19 | 1 | 0 | 2 | 1 | 0 | - | - |
| Patient 20 | 0 | 0 | 2 | 0 | 0 | - | - |
| Patient 21 | 1 | 0 | 6 | 1 | 0 | 1 | 0 |
| Patient 22 | 1 | 0 | 2 | 1 | 0 | - | - |
| Patient 23 | 1 | 0 | 6 | 0 | 0 | - | - |
| Patient 24 | 0 | 1 | - | - | - | 0 | 1 |
| Patient 25 | 1 | 0 | - | - | - | - | - |
| Patient 26 | 0 | 0 | - | - | - | 0 | 0 |
| Patient 27 | 0 | 0 | - | - | - | 0 | 0 |
| Patient 28 | 0 | 0 | - | - | - | - | - |
| Patient 29 | 0 | 0 | - | - | - | 0 | 0 |
| Patient 30 | 0 | 0 | 4 | 1 | 0 | 0 | 0 |
| Patient 31 | 0 | 1 | 16 | 0 | 1 | - | - |
| Patient 32 | 0 | 1 | 4 | 0 | 1 | 0 | 1 |
| Patient 33 | 0 | 1 | 10 | 0 | 1 | - | - |
| Patient 34 | 0 | 0 | 6 | 0 | 0 | 0 | 1 |
| Patient 35 | 1 | 0 | 52 | 1 | 0 | 1 | 0 |
| Patient 36 | 1 | 0 | 80 | 1 | 0 | - | - |
| Patient 37 | 1 | 0 | 30 | 1 | 0 | - | - |
| Patient 38 | 1 | 0 | 200 | 0 | 1 | - | - |
| Patient 39 | 1 | 0 | 20 | 0 | 0 | 1 | 0 |
| Patient 40 | 1 | 0 | 6 | 0 | 0 | - | - |
| Patient 41 | 0 | 0 | 4 | 0 | 0 | - | - |
| Patient 42 | 0 | 1 | 100 | 0 | 1 | - | - |
| Patient 43 | 0 | 0 | 80 | 0 | 0 | - | - |
| Patient 44 | 0 | 1 | 10 | 0 | 1 | 0 | 1 |
| Patient 45 | 1 | 0 | - | - | - | - | - |
| Patient 46 | 1 | 0 | - | - | - | - | - |
| Patient 47 | 0 | 0 | 2 | 0 | 0 | - | - |
| Patient 48 | 1 | 0 | 6 | 1 | 0 | - | - |
| Patient 49 | 1 | 0 | - | - | - | 1 | 0 |
| Patient 50 | 1 | 0 | - | - | - | - | - |
| Patient 51 | 0 | 0 | - | - | - | - | - |
| Patient 52 | 0 | 0 | 2 | 0 | 0 | - | - |
| Patient 53 | 0 | 1 | 12 | 0 | 1 | 0 | 1 |
| Patient 54 | 1 | 0 | 8 | 1 | 0 | - | - |
| Patient 55 | 0 | 1 | 2 | 0 | 1 | - | - |
| Patient 56 | 1 | 0 | 6 | 1 | 0 | - | - |
| Patient 57 | 0 | 0 | - | - | - | 1 | 0 |
| Patient 58 | 0 | 0 | - | - | - | - | - |
| Patient 59 | 1 | 0 | 14 | 1 | 0 | 1 | 0 |
| Patient 60 | 0 | 1 | - | - | - | - | - |
| Patient 61 | 1 | 0 | 60 | 0 | 0 | 1 | 0 |
| Patient 62 | 1 | 0 | 6 | 1 | 0 | - | - |
| Patient 63 | 0 | 1 | - | - | - | 0 | 1 |
| Patient 64 | 0 | 1 | - | - | - | 0 | 1 |
| Patient 65 | 1 | 0 | - | - | - | - | - |
| Patient 66 | 0 | 0 | - | - | - | - | - |
| Patient 67 | 0 | 0 | - | - | - | 0 | 1 |
| Patient 68 | 1 | 0 | - | - | - | 0 | 0 |
| Patient 69 | 0 | 0 | 12 | 0 | 0 | - | - |
| Patient 70 | 0 | 0 | 58 | 0 | 0 | - | - |
| Patient 71 | 0 | 0 | 116 | 0 | 0 | 0 | 0 |
